# Genome-wide association study elucidates the genetic architecture of manganese tolerance in *Brassica napus*

**DOI:** 10.1101/2024.03.27.586972

**Authors:** Harsh Raman, Zetao Bai, Brett McVittie, Sourav Mukherjee, Hugh D Goold, Yuanyuan Zhang, Nay Chi Khin, Yu Qiu, Shengyi Liu, Regine Delourme, Barry Pogson, Sureshkumar Balasubramanian, Rosy Raman

## Abstract

*Brassica napus* (canola) is a significant contributor to the world’s oil production and is cultivated across continents, yet acidic soils with Al^3+^ and Mn^2+^ toxicities limit its production. The genetic determinants underlying acidic soil tolerance in canola are unknown and require to be uncovered for canola breeding and production. Here, through comprehensive phenotyping, whole genome resequencing, and genome-wide association analysis, we identified three QTLs for tolerance to Mn^2+^ toxicity on chromosomes A09, C03, and C09. Allelism tests between four tolerance sources confirmed that at least one locus on A09 controls Mn^2+^ tolerance in *B. napus*. Integrated analysis of genomic and expression QTL and Mn^2+^ tolerance data reveals that *BnMTP8.A09,* in conjunction with *BnMATE.C03*, *BnMTP8.C04* and *BnMTP8.C08*, play a central role in conferring Mn^2+^ tolerance in *B. napus*. Gene expression analysis revealed a high correlation (*R*^2^ = 0.74) between Mn^2+^ tolerance and the *BnMTP8.A09* expression. Yeast complementation assays show that *BnMTP8.A09* can complement manganese-hypersensitive yeast mutant strain *PMR1*Δ and restore Mn^2+^ tolerance to wild-type levels. Inductively coupled plasma mass spectrometry revealed that Mn^2+^ tolerant accessions accumulate less Mn in the shoots compared to Mn^2+^ sensitives, suggesting that the *BnMTP8.A09* transporter likely sequesters Mn^2+^ into the tonoplast. Taken together, our research unveils the genetic architecture of Mn^2+^ tolerance and identifies *BnMTP8.A09* as a major gene imparting tolerance to Mn^2+^ toxicity in *B. napus*.

## Introduction

Soil acidity affects approximately 50% of the world’s arable land and limits the production of crops, especially in tropical and subtropical regions (Kochian, 1995). The projected effects of global climate change are likely to exacerbate Mn toxicity over the coming decades (Fernando and Lynch, 2015). At low pH (<5.5), exchangeable Al^3+^, Mn^2+^, and H^+^ ions get solubilized into a solution form, which causes toxicities to plants. With growing global food demands, increasing productivity from the marginal and problematic soil is essential. The relative importance of each ion toxicity varies across soils with different chemistries: Al^3+^ toxicity primarily inhibits root growth; while, Mn^2+^ toxicity causes interveinal and leaf margin chlorosis, brown and necrotic lesions, leaf cupping, and crinkling; with both result in reduced crop yield (Marschner, 1995, Foy, 1983, Bergmann, 1992). Mn^2+^ toxicity can also result in the inhibition of net photosynthesis assimilation, accumulation of reactive oxygen species, disruption of the activity of critical enzymes, and impairment of absorption, translocation, and utilization of essential nutrients for plant growth, including Ca^2+^, Fe^3+^, Zn^2+^, and Mg^2+^ (Horst, 1988, Bloom and Lancaster, 2018, LI, 2021). Extreme climatic conditions such as waterlogging with low redox potential, water deficit, and heat episodes can lead to excessive Mn^2+^ absorption and toxicity, affecting plant physiology and development across different soil types (Sparrow and Uren, 1987).

Surface soil acidity can be ameliorated by applying lime (CaCO_3_). However, it is challenging to incorporate lime in deeper layers to correct the subsoil acidity, and it takes several years before these soils become productive for commercial cropping. To cope with toxic levels of Mn^2+^ ions, plants have evolved several strategies, such as sequestration into subcellular compartments, activation of the antioxidant system, and regulation of the uptake, translocation, and distribution of Mn (Fecht-Christoffers et al., 2006, Peiter et al., 2007, Li et al., 2019). Several proteins play a role in the homeostasis and detoxification of Mn^2+^. These include various transporters such as Natural Resistance Associated Macrophage Protein (NRAMP1, NRAMP3, NRAMP4, NRAMP5), Iron Transporter (IRT1), ATP-binding cassette (ABC, multidrug resistance-associated proteins, MRP), iron-regulated transporters (IREG) and Cation/H^+^ Exchanger (CAX, CAX2, ECA1), Cation diffusion Facilitator (CDF, or metal tolerance protein, MTP), and P-type ATPase (ZIP) (Castaings et al., 2021, Li et al., 2019).

Natural variation for tolerance to Mn^2+^ toxicity is described in several plant species, including *Brassica napus* (Foy, 1984, Horiguchi, 1988, Khan and McNeilly, 1998, Basu et al., 2001, Schaaf et al., 2002, Kassem et al., 2004, Peiter et al., 2007, Mizuno et al., 2008, Pradeep et al., 2020, Wratten and Scott, 1979, Moroni et al., 2003, Delhaize et al., 2003, Raman et al., 2017). Studies have shown that *AtMTP11, BnMTP8.C04, BnMTP9.A07, ShMTP1,* and *NRAMP5* confer tolerance to Mn^2+^ tolerance in plants (Gu et al., 2022, Delhaize et al., 2003, Delhaize et al., 2007, Noor et al., 2023). However, the genetic basis of natural variation in tolerance to Mn^2+^ toxicity was not described in diverse *B. napus* germplasm.

*B. napus* (canola/rapeseed, 2*n* = 4*x* = 38, genome *A^n^A^n^C^n^C^n^*) is a widely grown critical crop of importance to agriculture and is sensitive to Mn^2+^ toxicity. It contributes approximately 12% of the global edible vegetable oil supply (FAO STAT, https://www.fao.org/faostat/) and protein for feedstock. In addition, canola oil accounts for 80%–85% of the renewable sources for biodiesel production (Tursi, 2019). To develop Mn^2+^ tolerant canola cultivars suitable for cultivation on acid soils, we have previously mapped a locus, *BnMTP8.A09,* for tolerance to Mn^2+^ toxicity (Raman et al., 2017) near the chromosome A09 orthologues of the *AtMTP8* transporter gene of *A. thaliana* (Delhaize et al., 2003). QTL mapping studies often capture only a slice of the genetic architecture of a trait because only alleles that differ between parental lines segregate (Holland, 2007). Understanding the genetic architecture of Mn^2+^ tolerance genes in genetically diverse germplasm provides insights into crop adaptation and assists the development of canola varieties.

Herein, we present the genetic architecture of Mn^2+^ tolerance in canola, which shows multiple loci contribute to tolerance to Mn^2+^ toxicity in diverse germplasm. In addition, we characterize and demonstrate that *the BnMTP8.A09* gene is a key candidate that explains a substantial proportion of variation in Mn^2+^ toxicity and is associated with tolerance Mn^2+^ toxicity under laboratory, glasshouse, and field conditions.

## Results

### Genotypic variation for Mn^2+^ tolerance in 326 *B. napus* accessions

For elucidation of the genetic architecture underlying tolerance to Mn^2+^ toxicity, we carried out six experiments (Method S1). To screen for Mn^2+^ tolerance we first scored symptoms of Mn^2+^ toxicity after five days of Mn^2+^ treatment in 415 accessions of *B. napus* (Table S1). Mn^2+^ toxicity was typically characterized by extensive chlorosis on the cotyledonary lobes (Figure 1A). Several accessions showed interveinal and marginal leaf chlorosis with some displaying cupping and necrotic spots on the leaf, suggestive of Mn^2+^ sensitivity. However, there were Mn^2+^ tolerant accessions which revealed no such visible symptoms or limited chlorosis (Figure 1B-F). We scored each accession for their Mn^2+^ tolerance, visibly on a scale of 1 to 5, While a majority of accessions (74%) were generally sensitive to Mn^2+^ with scores above 3, we observed significant variation among diverse accessions (Table S2, Figure 1G).

**Figure 1.**
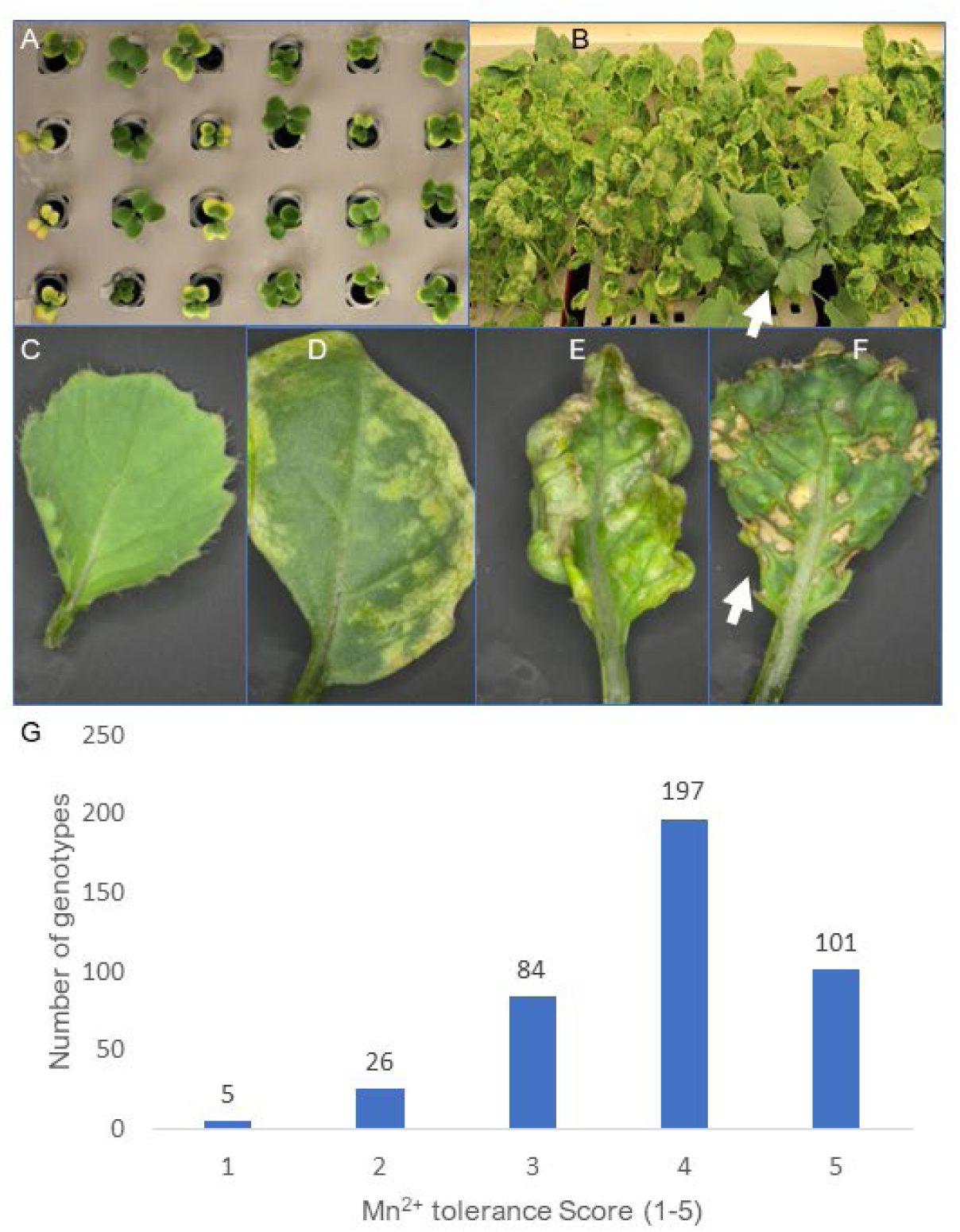
Critical symptoms and natural variation in tolerance to manganese (Mn^2+^) toxicity in *B. napus* accessions as evident in cotyledon and fully expanded leaves. (A) Mn^2+-^sensitive accessions show extensive chlorosis on cotyledons of young seedlings grown on 125 µM MnCl_2_. 4H_2_O on day 4 from germination, while Mn^2+^ tolerant accessions did not show such symptoms, as observed in Mn^2+^ sensitive accessions. (B) Mn^2+^ sensitive accessions show extensive chlorosis, curling and necrosis on mature leaves of seedlings (21 days after germination), while Mn^2+^ tolerant accessions show no such symptoms observed in Mn^2+^ sensitive accessions (marked with arrow). Stereomicroscope images showing normal leaf development in Mn^2+^ tolerant accession (C) and a range of toxicity symptoms (D: chlorosis; E: leaf curing and chlorosis F: dark brown leaf speckles on 3–4-week-old plants) in Mn^2+^ sensitive accessions. (G) Frequency distributions of Mn^2+^tolerance scores in the 326 *B. napus* accessions of GWAS panel. The tolerance scores are based on the extent of leaf chlorosis. Each line had four plants/replicate and replicated thrice (12 biological replicates).

### Genome-wide association analysis (GWAS) reveals genetic architecture for Mn^2+^ tolerance

To ascertain the genetic relatedness of the accessions, we carried out whole genome resequencing (WGR) at a moderate depth (4.69× to 99.59× with an average of 13.03×, Table S3). WGR provided 8,789,769 high-quality single nucleotide polymorphisms (SNPs) mapped to the reference genome of *B*. *napus* cv. Darmor-*bzh* v.4.1. Filtering-out variants with minor allele frequencies (MAF) <0.05 and missing rate >0.9 resulted in a total of 2,226,172 high-quality SNPs used for GWAS analysis, which averages to roughly one SNP marker per 20Kb of the canola genome. Population structure analysis revealed four genetically distinct clades: I, II, III, and IV in the GWAS population, consistent with their geographic origins (Figure 2A-C, Table S4). We calculated pairwise SNP linkage disequilibrium (LD) using 1,981,597 tagged SNP derived from haploblocks. The LD patterns were variable across the whole (*A^n^C^n^*) genome and within the *A^n^* and the *C^n^* subgenomes. Consistent with previous studies (Chalhoub et al., 2014), the LD decays faster in the *A^n^* subgenome than in the *C^n^* subgenome (Figure 2D).

**Figure 2:**
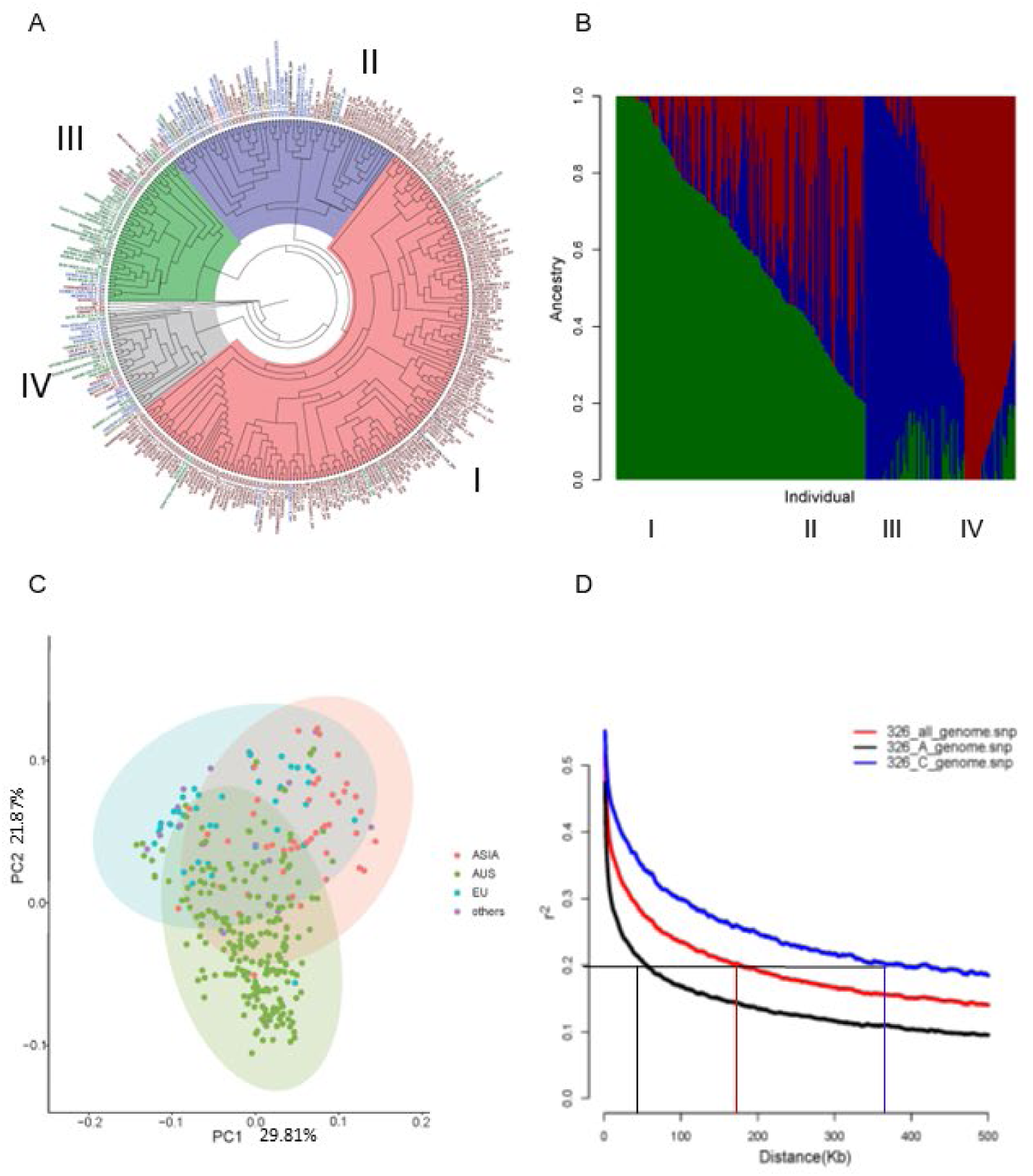
Genetic diversity, population structure and linkage disequilibrium in the AHGDS panel of *B. napus*. (A) Circular phylogenetic tree showing grouping (I-IV) of 326 accessions based on the neighbour-joining method. (B) population structure of 326 accessions revealed by the Bayesian method, STRUCTURE. (C) Principal component plots show three predominant clades of 326 accessions: Australia (I), Europe (II), Asia (III), and others (IV). (D) Genome and subgenome-wide LD decay plots. The horizontal black lines are the standard critical *R^2^* value, and the vertical red, black, and blue lines represent the AC, A and C subgenomes of *B. napus*.

Using Mn^2+^ toxicity phenotypes of the cotyledons and accounting for population structure and kinship coefficients, we identified 34 significant SNP associations (binned into three loci) for Mn^2+^ tolerance on chromosomes A09, C03, and C09 (Figure 3A, Figure S1B, Table S5). There was a good fit between observed and expected SNP associations (Figure 3B). Twenty-nine SNP associations (of 34 SNPs) were identified within the 300 kb genomic region on A09 (Table S5). Our results indicate that multiple loci contribute to tolerance to Mn^2+^ toxicity.

**Figure 3:**
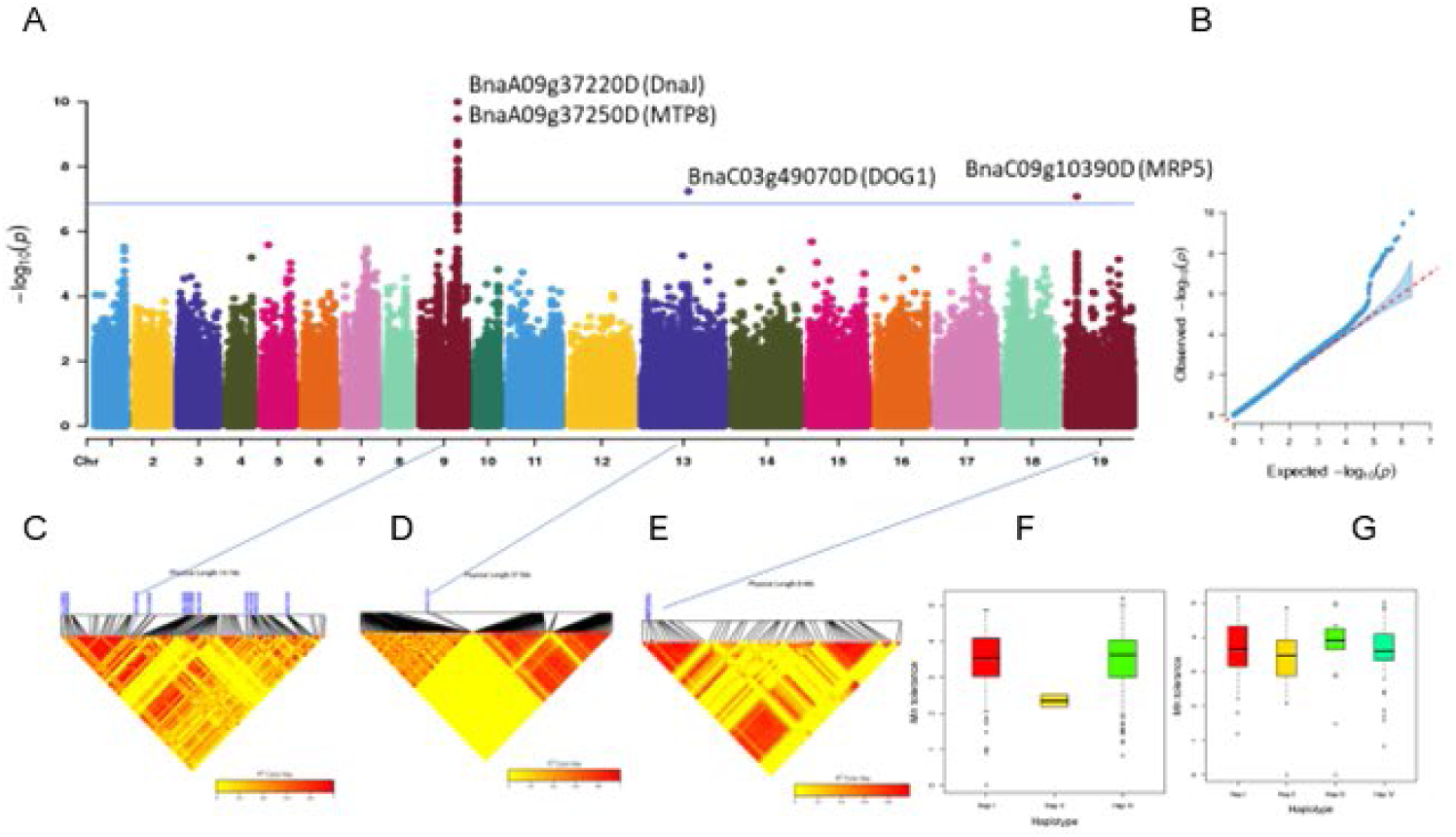
Natural variation in Mn^2+^ tolerance loci in *B. napus*. (A) Manhattan plots showing SNP associations for Mn^2+^ tolerance in a GWAS panel of *B. napus*. Three genomic regions (A09, C03 and C09) were associated with Mn^2+^ tolerance. Only QTL above the threshold– log(p-value) >2.5E^-8^ were included in this figure. The closest candidate gene (with suffix *Bna*) that maps near the highly significant SNP is also shown. (B) QQ plot showing a relationship between observed (y-axis) and expected (x-axis) LOD scores and a line of fit (red dashed line). Local linkage disequilibrium (LD) heatmap of the genomic region containing the most significant SNP associated with Mn^2+^ tolerance on chromosomes A09 (C), C03 (D), and C09 (E). The genomic region is based on the physical map position of the most significant SNP on the Darmor-*bzh* reference assembly ± 10kb. LD estimated as *r^2^* is shown in colour keys. Haplotype showing association with Mn^2+^ tolerance on chromosome A09 (F) and C09 (G). Box plots for Mn^2+^ tolerance grouped by alleles of the top SNP markers (maximum LOD scores) on A09, C03 and C09 chromosomes. The central bold line within the box represents the median; box edges indicate the upper and lower quartiles; whiskers show the 50% interquartile range, and points indicate outliers. Two-tailed two-sample Wilcoxon tests determined *P*-values.

### Validation and fine mapping of the GWAS loci using bi-parental populations

To verify the linkage between Mn^2+^ tolerance and SNP markers, we performed selective sweep analysis using *F_ST_* and ratio test, utilizing a cohort of 50 extreme accessions (25 tolerant accessions having mean score ≤ 2, group1 and 25 sensitive accessions with a mean score of ≥4, group 2) from the GWAS panel (Table S6). Although current breeding programs are not intentionally selecting for Mn^2+^ tolerance, our results showed that the A09 genomic region is subjected to passive selection under acid soil conditions (Figure S1A). Haplotype association analysis revealed that A09 haplotypes showed a statistically significant difference in Mn^2+^ tolerance (Figure 3C-G).

To validate the genetic linkage between Mn^2+^ tolerance and A09 genomic region, we generated two F_2_ populations derived from P3083 (China, tolerant to Mn^2+^) × ZY003 (China, sensitive to Mn^2+^), and Mutu (Japan, tolerant to Mn^2+^) × RSO 96 (sensitive to Mn^2+^), and tested them in hydroponic culture. Both populations showed approximately monogenic segregation for Mn^2+^ tolerance in a dominant manner (Table S7, Figure S2). QTL analysis revealed a single genomic region for Mn^2+^ tolerance on chromosome A09 (Figure S3). We compared the physical positions of SNPs associated with Mn^2+^ tolerance with those identified in the earlier study from Darmor-*bzh*/Yudal (Raman et al., 2017). We found that one genomic region on chromosome A09 (26.36 Mb to 27.16Mb) is shared across the GWAS, F_2_, and DH (Darmor-*bzh*/Yudal) panels (Table S5), suggesting of common allelic variation at this locus for Mn^2+^ tolerance in *B. napus*. To test whether Mutu and Darmor-*bzh*, earlier described sources of Mn^2+^ tolerance (Moroni et al 2003, Raman et al 2017) also harbour the same gene(s), we made crosses between three sources of tolerance. There was no segregation among F_2_ progenies derived from Darmor-*bzh* (France, tolerant to Mn^2+^) × Mutu (Japan, tolerant to Mn^2+^) and Darmor-*bzh* (tolerant to Mn^2+^) × Jet Neuf (France, tolerant to Mn^2+^) TableS7, Figure S2). These results suggest that parental lines Darmor-*bzh*, Jet-Neuf, and Mutu have the same or similar alleles that control Mn^2+^ tolerance. This result also corroborates that Darmor-*bzh* and Jet-Neuf share a common ancestry.

### Candidate genes underlying significant loci contributing to Mn^2+^ tolerance

We searched candidate genes based on LD with significantly associated SNPs (Table S5) and found 19 genes that were located within the 7.8 kb regions of the GWAS-SNPs on A09, C03, and C09 chromosomes (Figure S1B-D, Table S8), including Metal Tolerance Protein 8 (MTP8, designated as *BnMTP8* in *B. napus*), which has previously been shown to confer natural variation in Mn^2+^ tolerance in Arabidopsis (Delhaize et al., 2003) making it an obvious candidate for further investigation. Protein-protein-interaction network using the STRING (search tool for recurring instances of neighbouring genes) database (Version 11.0, http://string-db.org/) also revealed that candidate genes prioritized in this study are related to cation transport and intercellular homeostasis. Candidate genes include several Zn transporters (*ZAT*, AT2G04620, AT3G12100, *MTPB1*, AT1G51610, *MTPA2*), high-affinity Mn^2+^ transporter involved in Mn, Fe, Cd and Co acquisition (NRAMP1), vacuolar transporter involved in intercellular metal (Fe, Mn, Cd and Co) homeostasis, IREG2, (IRON REGULATED 2) encoding FPN2, a tonoplast localized Ni transport protein network with MTP8, and TMN1 (Transmembrane 9) gene which interacts with proteins involved in Golgi transport complex-related vascular transport, inter-cellular protein transport, and transfer from ER via Golgi (Figure S1E-G).

We sequenced the full length of *BnMTP8* alleles (1966 bp) from the parental lines of the Darmor-*bzh*/Yudal DH population using A09 sequence-specific primers, which revealed 12 polymorphic SNPs and InDELs (Table S9, Figure 4A). In addition, we characterised sequence variants in *BnMTP8* from a dataset of 2,289 *B. napus* sequenced accessions (http://yanglab.hzau.edu.cn/BnIR). These data revealed that the *BnMTP8.A09* downstream sequence had the most sequence variants (55.6%) consistent with its association suggestive of that being the candidate gene (Figure 4B-C).

**Figure 4:**
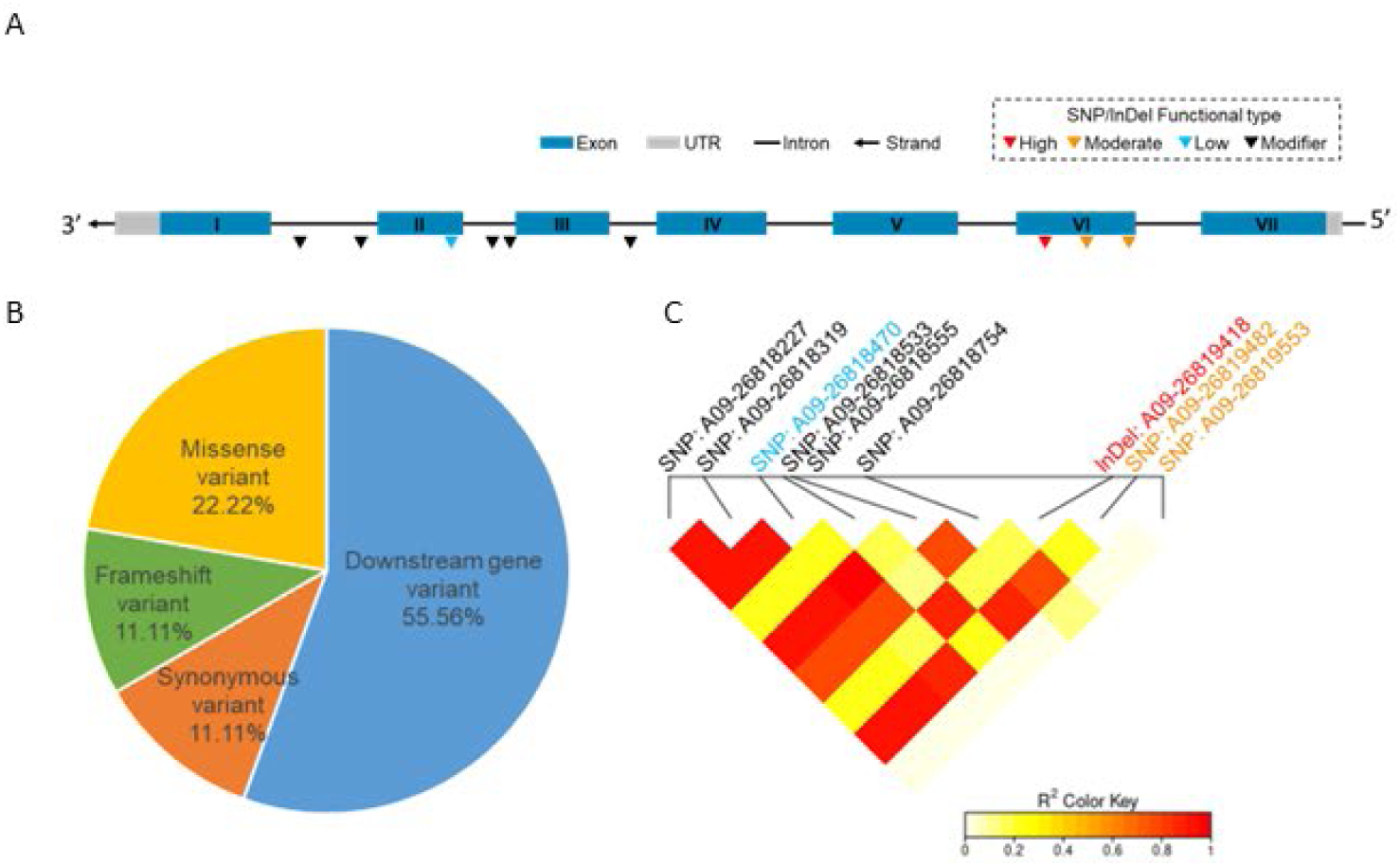
Structural variants and their distribution in *BnMTP8.A09* (*BnaA09g37250D)* gene. (A) Schematic representation of the *BnaA09g37250D* (*BnMTP8.A09,* 1966 bp) gene encoding a cation diffusion facilitator protein showing seven exons and six introns in black and grey colour, respectively. The inverted triangle symbols indicate natural population SNPs/InDels located on *BnaA09g37250D*. The colours of inverted triangle symbols indicate different variations of functional types. (See detailed information in Table S4B). B: Annotation and the proportion of SNPs and InDels of *BnaA09g37250D* in 2289 accessions *of B. napus.* C: Linkage disequilibrium heatmap of population SNPs/InDels on *BnaA09g37250D*.

### Gene expression variation in *BnMTP8.A09* explains 74% of natural variation in Mn^2+^ tolerance

To assess whether *BnMTP8.A09* is the major locus associated with Mn**^2+^** tolerance in *B. napus*, we performed a selective DNA genotyping of 20 DH lines of Darmor-*bzh*/Yudal segregating for Mn^2+^ tolerance by Sanger sequencing. We found a complete linkage between *BnMTP8.A09* alleles and Mn**^2+^** tolerance (Figure S4A). To assess whether the expression level of the *BnMTP8* gene correlates with phenotypic difference in Mn^2+^ toxicity, we compared the expression levels of *BnMTP8.A09* in extreme accessions which were categorised as sensitive (S) or tolerant (T) to Mn^2+^ toxicity (Table S10). There are six copies of the *BnMTP8*, located on the homoeologous regions on the *A^n^* and *C^n^* subgenomes (A04/C04, A07/C06 and A09/C08) differ only by a few SNPs (Figure S4B, Table S9). We designed multiple primers with a 3’ mismatch to specifically amplify and quantify the *MTP8* sequence on chromosome A09 (gene ID: BnaA09g37250D in Darmor-*bzh* v4.1 reference, Table S11). The amplified products were sequence verified to ensure that PCR amplification is specific to only *BnMTP8.A09*. Our analysis revealed a striking correlation between sensitive and tolerant lines, with the expression levels being visibly high in tolerant lines compared to sensitive ones (Figure 5A, Figure S5). Quantitative reverse-transcription PCR analysis revealed that expression levels of *BnMTP8.A09* could explain up to 74% variance in the Mn^2+^ tolerance phenotype (*R^2^*=0.74, Nominal Logistic regression with expression level as a factor and phenotype (sensitive/tolerant) as a response, *p* < 0.0001). In addition, among the tolerant varieties, there was a marginal yet significant increase in *BnMTP8.A09* expression upon Mn^2+^ treatment (Fig. 5B). Based on this striking correlation, we conclude that Mn^2+^ dependent transcriptional control of *BnMTP8.A09* expression levels accounts for most of the variation in Mn^2+^ toxicity in *B. napus* accessions.

**Figure 5:**
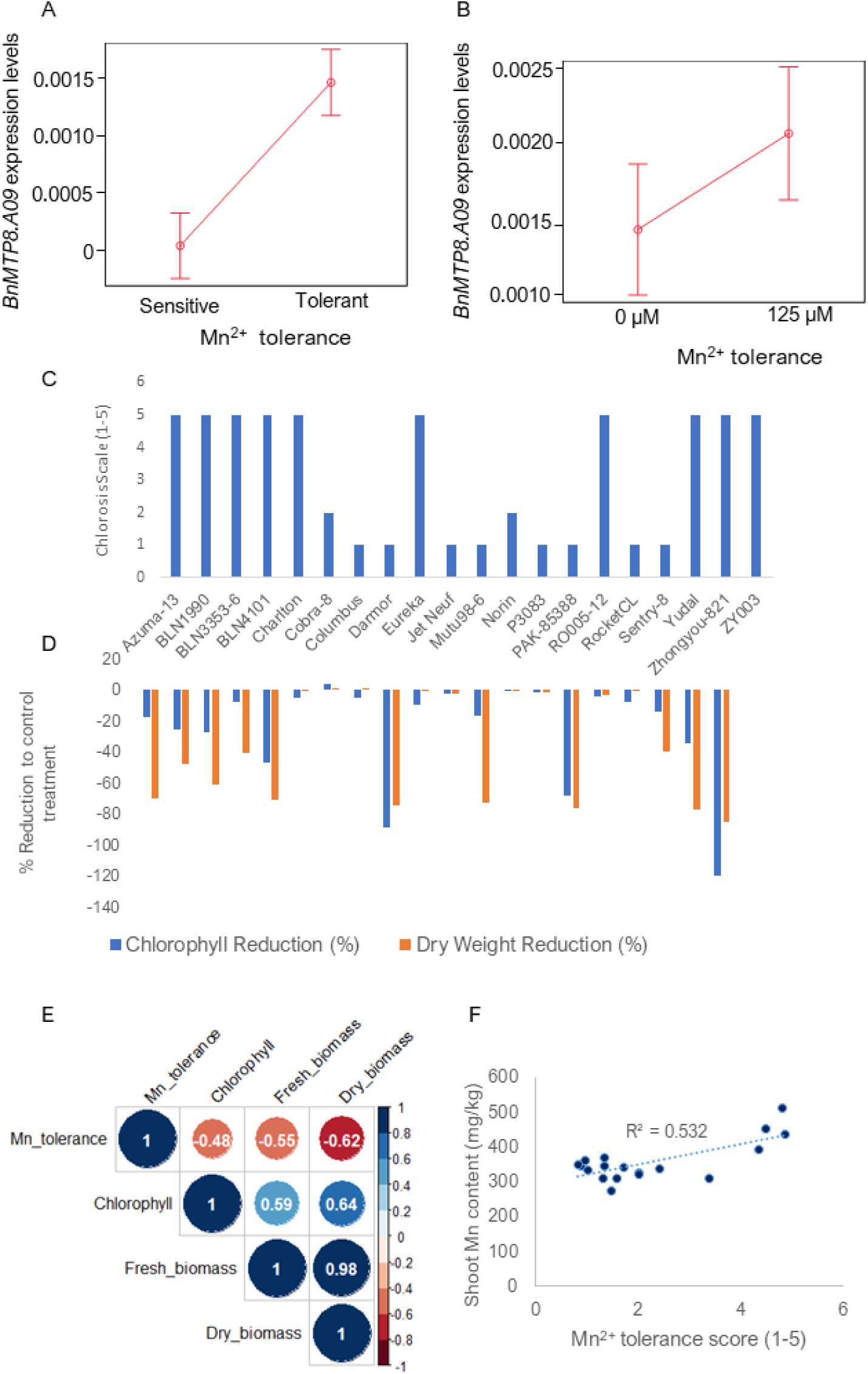
MTP8 functions in Mn^2+^ tolerance in *B. napus*. (A) Relative expression levels of *MTP8* in Mn^2+^ tolerant (T, n =) and sensitive (S) lines (n=) and (B) of 20 diverse accessions of *B. napus* after 6 days of stress in the 125µM MnCl_2_.4H_2_O and the corresponding control (9µM of MnCl_2_. 4H_2_O, –ve Mn^2+^ treatment). (C) Genetic variation for Mn^2+^ tolerance among 20 diverse accessions of *B. napus*. Chlorosis was measured as 1 (tolerant) to 5 (sensitive) scale. D: Per cent reduction in chlorophyll content and shoot weights of 20 accessions that show the extreme phenotypes for Mn^2+^ tolerance and sensitivity measured after 3-4 weeks in control nutrient solution (-ve Mn^2+^ treatment) and with 125 µM of MnCl_2_. 4H_2_O (C). Chlorophyll content was measured with SPAD. Above-ground shoot biomass was measured on a fresh and dry weight basis. (D) Per cent reduction to control treatment was calculated as the Trait value of Mn^2+^ plus treatment minus trait value of the control treatment)/trait value to control treatment x 100. The means of 12 biological replicates were plotted in the R package. (E): Phenotypic correlation between Mn^2+^ tolerance, chlorophyll content, and fresh and dry biomass of 20 accessions. (F): Correlation between Mn**^2+^** tolerance scores and shoot Mn^2+^ content.

We further raised the same 20 diverse accessions differing in *BnMTP8.A09* expression (Figure 5C) with and without Mn^2+^ treatments in controlled environment conditions (22°C, 50% humidity, 16/8 h). As expected, tolerant lines showing high *BnMTP8.A09* expression lines did not show critical symptoms of Mn^2+^ toxicity and accumulated 5.04× higher biomass than Mn^2^ sensitive and low *BnMTP8.A09* expression lines. Low *BnMTP8.A09* expression lines carrying sensitive alleles for Mn^2+^ tolerance showed reduced SPAD values, a proxy for chlorophyll content of the expanded leaf, and shoot biomass in high Mn^2+^ concentration (125µm MnCl_2_) compared to tolerant lines with high *BnMTP8.A09* expression (Figure 5C-D). Mn^2+^ tolerance positively correlated with fresh and dry shoot weights (Figure 5E-F). The chlorophyll (SPAD values) had a negative relationship with Mn^2+^ tolerance (Figure 5E). Chlorophyll is essential for net photosynthesis assimilation, which could affect biomass production. In previous studies, high biomass was positively related to seed yield in *B. napus* (Raman et al., 2016, Raman et al., 2020).

To directly assess whether these lines differ in Mn^2+^ accumulation, we performed inductively coupled plasma mass spectrometry (ICP-MS) analysis, which revealed that *B. napus* lines sensitive to Mn^2+^ toxicity accumulated more manganese than Mn^2+^ tolerant lines; shoot Mn content and Mn^2+^ tolerance score were positively correlated (*r* = 0.75, Figure 5F). Furthermore, Mn concentration negatively correlated (*r* = 0.5) with Fe and positively correlated with Ca (*r* = 0.8) accumulation (Figure S6), suggesting that Mn^2+^ toxicity could be related to Fe deficiency in the acidic soils.

### *BnMTP8.A09* is a bonafide Mn^2+^ transporter

To confirm the contribution of the *BnMTP8.A09* gene in natural variation in Mn^2+^ tolerance, we performed a complementation assay using the Mn-hypersensitive yeast strain *pmr1*Δ. *Pmr1* is a P-type ATPase responsible for transporting Ca and Mn into the Golgi apparatus, a major pathway for the cellular detoxification of manganese (Antebi and Fink, 1992). Yeast assays revealed that the *BnMTP8.A09* gene imparts the Mn^2+^ tolerance in the yeast strain *pmr1*Δ, which tolerated elevated Mn levels (50mM, i.e., 400× dose), which enabled discrimination of Mn^2+^ tolerant lines from sensitive ones in nutrient solution (Figure 6). These findings suggest that *BnMTP8.A09* is a bonafide Mn^2+^ transporter.

**Figure 6:**
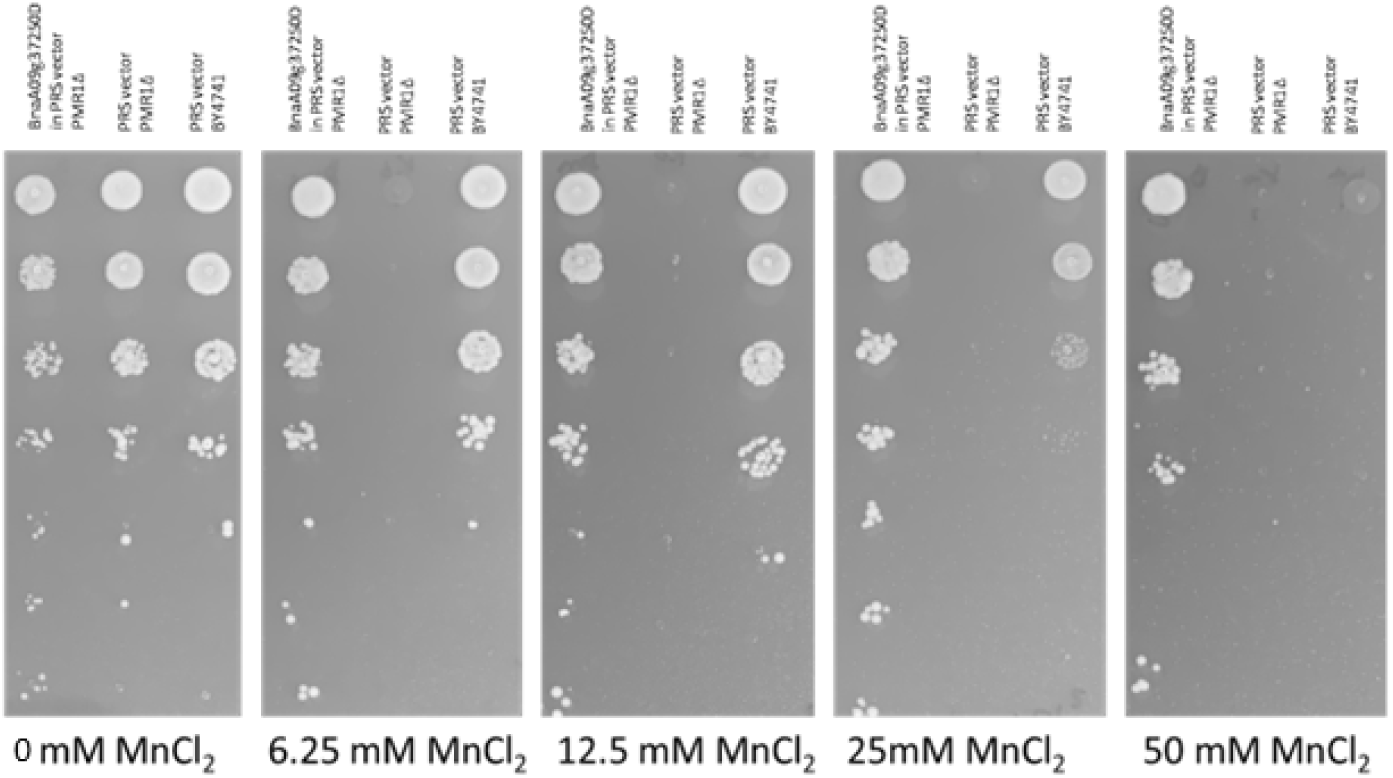
Effect of the *B. napus MTP8* expression on tolerance to Mn^2+^ toxicity. Yeast cells (Mn^2+^ hypersensitive yeast mutant *pmr1*Δ) carrying empty PRS vector, *BnMTP8* gene (BnA09g37250D in PRS vector) were spotted on the yeast medium (pH 4.4) without MnCl_2_ (control) and with MnCl_2_ concentrations (6.25, 12.5, 25 and 50 mM). The plates were incubated for 48 h and photographed.

### Expression QTL analysis revealed the central role of *BnMTP8.A09* in the genetic architecture of Mn^2+^ tolerance

To investigate what regulates the expression of *BnMTP8.A09*, which explained close to 74% of the variation in Mn^2+^ tolerance, we carried out the expression quantitative trait loci (eQTL) mapping of *BnMTP8.A09* in the *B. napus* population using RNA sequencing data from seedling leaves of 154 *B. napus* accessions (Figure S7A). Four eQTL of *BnMTP8.A09* were detected on chromosomes A09, C03, C04 and C08 (Figure 7A) and three of them colocalized with Mn^2+^ tolerance-related loci (Figure 3A, 7A). Based on the physical distance between eQTL and the target gene, we found the eQTL on A09 is *cis*-eQTL (< 1Mb) of *BnMTP8.A09. BnMTP8.A09* showed a higher expression among different homologues (Figure S7B) and is *cis*-regulated by SNPs of its flanking sequence. These findings strongly support our thesis that *BnMTP8.A09* is the target causal gene in the QTL of Mn^2+^ tolerance. Interestingly, we found three *trans*-eQTL of the *BnMTP8* homologues and two of them were on C04 and C08 (Figure 7B); although, there was no significant GWAS SNP within the two gene loci probably due to their expression dosage compensation effect on *BnMTP8.A09*. These results indicated that *BnMTP8.A09* has the most important effect among *MTP8* homologs. For the colocalized eQTL and QTL on C03, we found a candidate gene, Fe homeostasis-related FERRIC REDUCTASE DEFECTIVE3 (*BnaC03g49020D*, *BnFRD3.C03/BnMATE.C03*), a member of the MATE gene family that is also involved in cross-talk with Zn tolerance (Rogers and Guerinot, 2002, Pineau et al., 2012), contribute to Mn^2+^ tolerance and also could have genetic interaction with *MTP8* involved in Mn^2+^ tolerance network. The QTL of Mn^2+^ tolerance on A09 also showed epistatic effects with the QTL on C03 and C09 (Table S12). Nevertheless, these results indicated that *BnMTP8.A09* plays a central role in Mn^2+^ tolerance in *B. napus*. Next, we asked which of the specific polymorphism(s) could explain most of the variation in *BnMTP8.A09* of *B. napus* 2,280 accessions. We found that only an InDel(Ref/Alt = AT/A), in nine Asian and European accessions (Figure 7C, D), showed a significant association with *BnMTP8.A09* (BnaA09g37250D) expression in 289 *B. napus* accessions (Figure 7D). Interestingly, the same InDel was previously found to be associated with seed oil content (Figure S7E), suggestive of pleiotropy (Yang, 2023)

**Figure 7:**
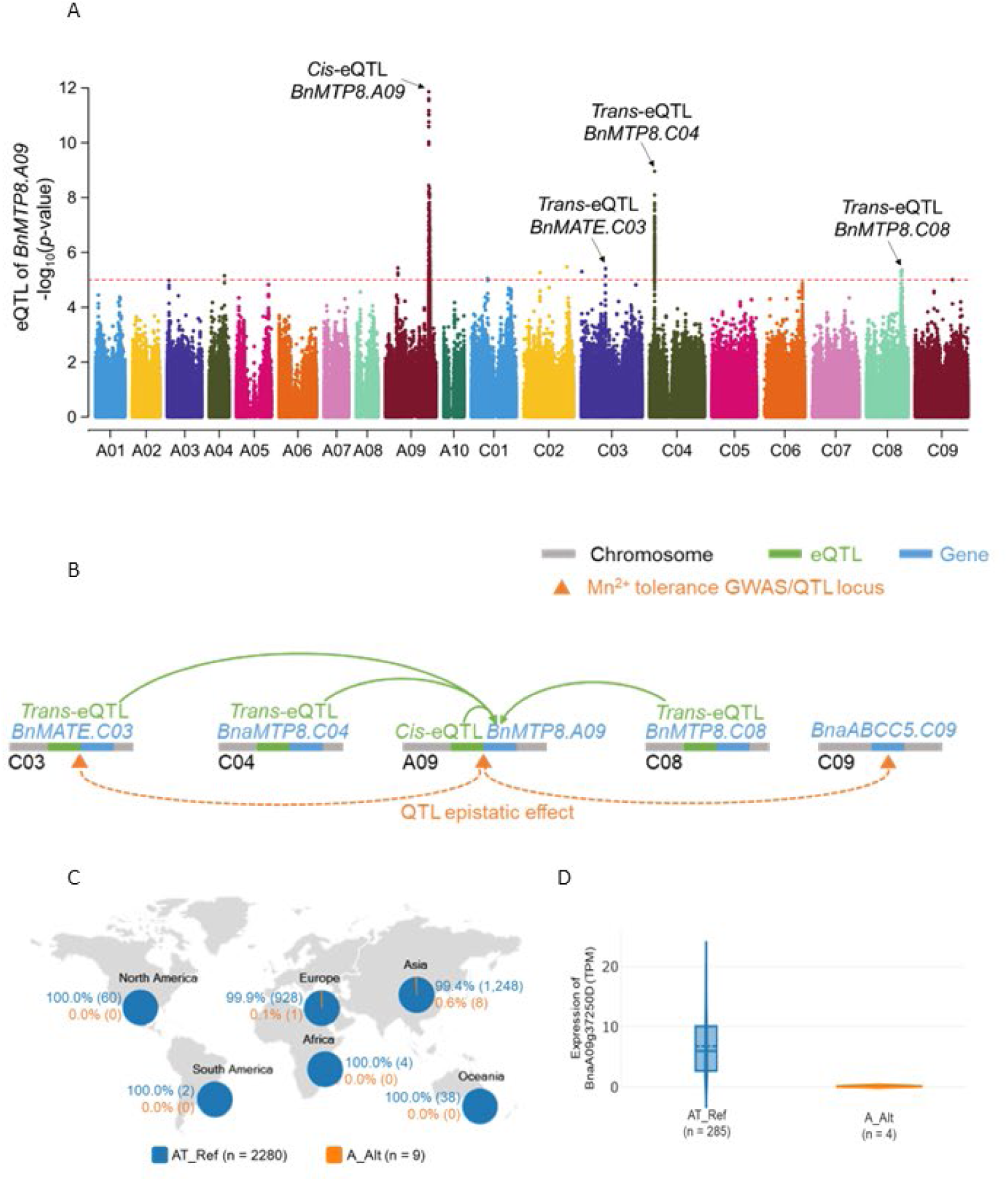
Expression QTL (eQTL) analysis revealing the genetic architecture of Mn^2+^ tolerance in *B. napus*. (A) Manhattan plot of eQTL for *BnMTP8.A09* in x *B. napus* accessions. Each point represents an SNP/InDel in the *B. napus* GWAS population. The candidate-expressed genes (*BnMATE.C03*, *BnMTP8.C04* and *BnMTP8.C08*) in *trans*-eQTL which could influence the expression of *BnMTP8.A09* were marked. (B): Regulatory network of *BnMTP8.A09* mediated the response to Mn^2+^ tolerance in *B. napus*. One *cis*-eQTL and three *trans*-eQTL colocalized with Mn^2+^ tolerance QTL on Chromosomes A09, C03, C04 and C08 indicated that *BnMTP8.A09* play an important role in Mn^2+^ tolerance in *B. napus*. The epistatic effects of the QTL between A09 and C03/C04/C08/C09 suggest the complex regulatory mechanism of Mn^2+^ tolerance in *B. napus*. The green line (eQTL) next to the short blue line (candidate gene) indicates the corresponding candidate gene located in the eQTL region. (C) Geographic distribution of 2,289 *B. napus* accessions with the allelic variation of target InDel on *BnaA09g37250D*. Each pie indicates the proportion of *B. napus* accessions with the allelic variation of target InDel (InDel: A09-26819418) located in *BnaA09g37250D* in six continents, respectively. The proportion and the number next to each pie indicate the proportion and total number of accessions with target SNP alleles. Blue and orange indicate the two allelic variations. “n” is the total number of accessions with the allelic variation. (D): The violin plot reveals the difference in the gene expression level of *BnaA09g37250D* between the population allelic variation of target InDel (InDel: A09-26819418). Box shows the median and interquartile range values. “*n*” is the accession number for statistics.

### *BnMTP8.A09* alleles confer tolerance to Mn2^+^ toxicity under field conditions and are under purifying selection

To assess whether allelic variation in *BnMTP8.A09* indeed makes a difference to plants, we tested 175 doubled haploid (DH) lines from the Darmor-*bzh*/Yudal (DY) population with natural alleles that segregate for Mn^2+^ tolerance, along with parental lines of DY population and 15 controls in acid soil conditions at Mangoplah, NSW, Australia (2022, soil pH: 4.5, Mn = 198µM, Figure S8A). We observed critical symptoms for Mn^2+^ toxicity, manifested as leaf chlorosis, were visible at physiological maturity (BBCH GS89) when plots were waterlogged due to excessive rainfall and subjected to high temperatures (Figure S8B), though conditions (warm and dry winter and early spring) at earlier stages were not favourable to assess Mn^2+^ toxicity in the cotyledon stage. However, symptoms of Mn^2+^ toxicity were apparent in reshoots and plants regenerated from pre-harvest shattered seeds in the plots (Figure 8). We took an opportunistic approach, scored symptoms of Mn^2+^ toxicity (leaf chlorosis) on re-shooted plants, and related them with Mn^2+^ tolerance scores from the nutrient screening experiment. Interestingly, cotyledon chlorosis scores of DH lines assessed in nutrient culture (Raman et al., 2017) were positively correlated with leaf chlorosis in the field (*r* = 0.45, Figure S6A). To verify the results, we selected 20 lines from the DYDH population that had consistent scores for Mn^2+^ tolerance in nutrient solution and field conditions and had contrasting marker alleles associated with Mn^2+^ tolerance at *BnMn*^2+^*.A09* locus (Raman et al., 2017). These lines were evaluated in acidic soil having high Mn^2+^ (pH 4.5), collected from Mangoplah under glasshouse conditions at Wagga Wagga. Test lines showed differences in tolerance to Mn^2+^ toxicity and a positive correlation (*r* = 0.9) with cotyledon scores in nutrient culture. These findings conclusively demonstrated that *BnMTP8.A09* indeed confers tolerance to Mn^2+^ toxicity not only in lab conditions but also in field conditions.

**Figure 8:**
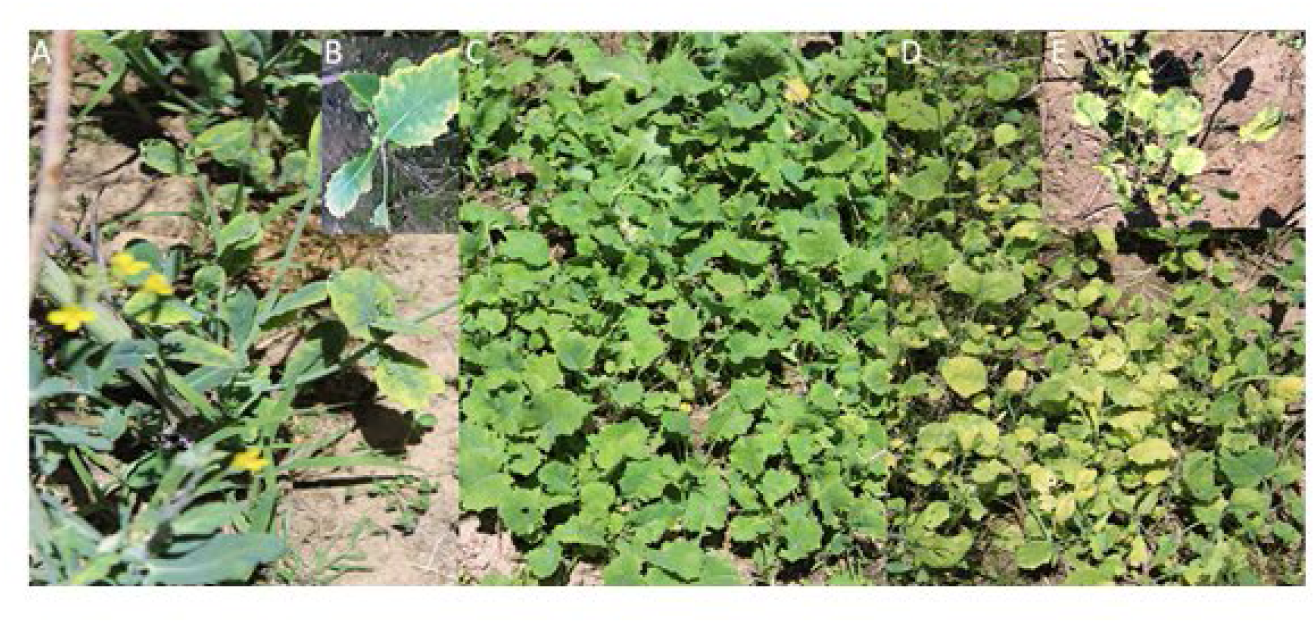
*Brassica napus* showing genetic variation for tolerance to Mn^2+^ toxicity under field conditions. A-B: Reshooted plant showing symptoms of leaf chlorosis; Shattered seeds in plots showing leaf chlorosis in Mn^2+^ sensitive (C-D, *Bnmtp8.A09*) and tolerant (E, *BnMTP8.A09*) doubled haploid lines of Darmor-*bzh*/Yudal population grown on acidic soil (pH 4.5) at Mangoplah NSW, Australia.

To assess whether *BnMTP8.A09* alleles may have undergone selection during crop domestication, we determined nonsynonymous sites (*Ka*)/synonymous sites (*Ks*) of *BnMTP8.A09*. Our results suggested that *BnMTP8.A09* is under purifying selection (p=0) irrespective of *B. napus* ecotypes (Figure S9). These findings suggest conservation of gene function across spring, winter and semi-winter types due to adaptation to environments with Mn^2+^ toxicity.

## Discussion

### Natural variation in Mn^2+^ tolerance in *B. napus*

*B. napus* was domesticated and selected for consumer preferences in Southern Europe, where carbonate-rich soils are highly prevalent (Gómez-Campo and Prakash, 1999, Prakash et al., 2011). In this study, we assessed a diversity panel representing spring, semi-winter and winter ecotypes and identified only five (1.2%) Mn^2+^ tolerant *B. napus* accessions (which had the least chlorosis scores of ≤1) of spring-type from Asia (Pakistan, Japan), and winter-type from Europe (France, Germany, Table S1-2). These results suggest that populations of *B. napus* had progressed adaptation mechanisms to minimize the harmful effects of toxic ions in acid soils by natural selection and/or by passive selection by breeders.

### *BnMTP8.A09* controls natural variation for Mn^2+^ tolerance in *B. napus*

Through GWAS, selective sweep and eQTL analyses, we uncovered 3 to 4 genomic regions for Mn^2+^ tolerance (Figures 3, 7, S1), suggesting that genome-wide approaches are suitable for revealing the architecture of Mn^2+^ tolerance. One of the highly significant association peaks was located close to the *BnMn^2+^.A09* locus, previously identified by QTL analysis in the *B. napus* DH population (Raman et al., 2017). However, no QTL/eQTL for Mn^2+^ tolerance on C03 and C09 chromosomes were identified in earlier studies. The *trans*-eQTL associated with Mn*^2+^* tolerance on chromosomes A09 and C03 (Figure 7) were mapped within the LD of GWAS-SNPs, suggesting that both genes (*BnMTP8.A09* and *BnMATE.C03*) are likely coregulated. Future studies need to validate those new genomic regions for their contribution to Mn*^2+^* tolerance.

We identified several highly significant SNP associations that were located near the transporter genes (Table S8). These transporters are implicated in metal ion transport in different plant species (Leung et al., 2019, De Caroli et al., 2020, Chu et al., 2017, Tsunemitsu et al., 2018). For example, MTP8 is a tonoplast localized member of the CDF and functions in roots as an Mn^2+^ transporter. In Arabidopsis, it transports Mn into root vacuoles of iron-deficient plants, thereby inhibiting iron deficiency-induced (ferric) chlorosis (Eroglu et al., 2016). MATE transporters are implicated in turgor-regulating chloride channels and xenobiotic detoxification by transmembrane export across the plasma membrane; this gene confers Al^3+^ tolerance by mediating citrate efflux from root cells in wheat, barley, sorghum and Arabidopsis (Upadhyay et al., 2019, Zhang et al., 2017) to chelate Al^3+^. The ABCC5 (C09) encodes a High-Affinity Inositol Hexakisphosphate Transporter. It plays a role in the signalling of guard cells, phytate storage, cellular potassium ion homeostasis, and response to salt stress (Lemtiri-Chlieh et al., 2000, Andolfo et al., 2015). ABC transporters are responsible for detoxifying many compounds from the cytoplasm (Klein et al., 2006).

### *BnMTP8.A09* imparts Mn^2+^ tolerance in canola and yeast

In this study, we show for the first time the functionality of *BnMTP8.A09* that underlies natural variation in Mn^2+^ tolerance in *B. napus* yeast complementation assay in a strain lacking the Golgi-mediated cytoplasmic efflux carrier PMR1 (Dürr et al., 1998), and phenotypic expression using natural alleles under controlled environment and field conditions. *MTP8* homologs have been shown to enhance tolerance to Mn^2+^ toxicity, Mn sequestration and plant growth in different species (Mills et al., 2008, Eroglu et al., 2016, Delhaize et al., 2003, Chen et al., 2013). For example, *AtMTP8* is reported to alleviate the antagonistic interference of Mn^2+^ with Fe^2+^ by loading Mn^2+^ into the root vacuole at high pH conditions in Arabidopsis (Eroglu et al., 2016, Farthing et al., 2023). In *B. napus*, three homologues; *BnMTP3*, *BnMTP8* (BnaCnng31720D on chromosome C04/*BnMTP8.C04*, AT3G58060D) and *BnMTP9.A07* (BnaA07g34970D, AT1G79520D) are shown to load Mn^2+^ into vesicles for subsequent delivery to the vacuole or secretion into extracellular spaces and maintain Mn^2+ h^omeostasi^s^ in the roots and shoots (Gu et al., 2022, Gu et al., 2021). In this study, ICP-MS analysis also suggested that Mn^2+^ tolerant accessions do not accumulate higher Mn in the shoots than Mn^2+^ sensitive lines (Figure 7D); therefore, tolerant accession could compartmentalize and sequestrate Mn^2+^ into the vacuole.

### Validation of high throughput method for screening germplasm for Mn^2+^ tolerance

Evaluating natural or transgenic *B. napus* lines under field conditions is challenging, as the phenotypic expression of Mn^2+^ toxicity depends on soil pH and weather conditions. Our results are consistent with literature that suggests warm and waterlogged conditions favour the expression of Mn^2+^ toxicity genes in acidic soil containing high exchangeable Mn^2+^. In addition, the critical symptoms of tolerance for Mn^2+^ toxicity and symptoms were variable across the field plots. It is emphasized that Mn^2+^ toxicity is not only a limitation in the acidic soil; the availability and/or unavailability of other ions, such as Al^3+^, H^+^, Fe^2+^, and Ca^2+^, could compromise the expression of Mn^2+^ tolerance under field conditions.

We rated the plots that showed no symptoms of Mn^2+^ toxicity as tolerant, while plots showing >10% of plants were scored as Mn^2+^ sensitive. Therefore, this criterion was subjected to ascertainment bias. Our data showed that hydroponic-based screening is more robust and reliable than field and glasshouse screening of large breeding germplasm that requires a large volume of soil with desired characteristics. Many plants can be screened in less than 10 days in a hydroponic system, providing highly reliable phenotyping for Mn^2+^ tolerance. High throughput phenotyping, in conjunction with molecular markers, could provide an efficient pipeline for tracking Mn^2+^ tolerance alleles in the *B. napus* breeding program. The resources developed herein would enhance selection efficiency in the breeding programs.

## Conclusion

Multiple lines of evidence support the conclusion that *BnMTP8.A09*, in conjunction with *BnMATE.C03*, *BnMTP8.C04* and *BnMTP8.C08*, play a significant role in conferring Mn^2+^ tolerance in *B. napus*. This includes GWAS, eQTL, QTL, gene expression profiling, yeast complementation, doubled haploids, Mn uptake analyses, with a range of phenotypic assays performed in laboratories, controlled environments, and field trials. Collectively, all of these indicate that expression of *BnMTP8.A09* is a major effect QTL for Mn tolerance in *B. napus*, such that gene expression variation in *BnMTP8.A09* explains 74% of natural variation in Mn^2+^ tolerance.

## Materials and Methods

### Plant materials

To investigate the genetic architecture of loci controlling tolerance to Mn^2+^ toxicity, we carried out six experiments in laboratory, glasshouse and field conditions (Method S1). The GWAS panel consisted of 415 *B. napus* spring, semi-winter and winter accessions representing Australian, Asian, North American and European breeding programs (Experiment 1).

To test the genetic inheritance, chromosomal locations and relevance of GWAS associations in the *B. napus* breeding programs, we generated three F_2_ intercross populations derived from P3083 (Chinese cultivar) × ZY003 (Chinese cultivar), Darmor-*bzh* (French cultivar) × Mutu (Japanese cultivar), and Darmor-*bzh* × Jet Neuf (French cultivar) (Experiment 2-3; Table 1). The relationship between Mn^2+^ tolerance and *BnMTP8* expression was established using a subset of GWAS (Experiment 4). For field validation of Mn^2+^ tolerance under field conditions, 175 doubled haploid (DH) population and its parental lines: Darmor-*bzh* and Yudal (Korean cultivar) population (Pilet et al., 1998, Raman et al., 2017) plus 15 controls (Table S1), complemented with a glasshouse bioassay (Experiment 5).

### Phenotypic evaluation for tolerance to Mn^2+^ toxicity

Genetic variation of *B. napus* accessions for tolerance to Mn^2+^ toxicity was assessed in different environments (Table S1). Response of *B. napus* GWAS and F_2_ lines to Mn^2+^ tolerance was assessed initially in a nutrient solution supplemented with 125μM of manganese tetrahydrate (MnCl_2_⋅4H_2_O) as described previously (Raman et al., 2017). Statistical valid experiment designs were followed and detailed (Methods S2). After 96 h of Mn^2+^ treatment, the critical symptoms of Mn^2+^ toxicity as the extent of chlorosis on cotyledonary lobes were scored quantitatively as “1” to “5” as described earlier (Raman et ^a^l., 201^7^). After scoring each F_2_ population and the parental lines for Mn^2+^ tolerance, 2 to 3-week-old seedlings from each population were transplanted in plastic pots to raise F_2:3_ progenies. Plants were selfed to ensure purity and avoid cross-pollination for different phenotyping and genotyping experiments. Each F_2:3_ line was assessed for Mn^2+^ tolerance as described above.

### Evaluation of lines for field and glasshouse performance

A total of 192 lines, including 175 from DH population derived from Darmor-*bzh* and Yudal, two parental lines (Darmor-*bzh* and Yudal) and 15 controls were evaluated to test their performance on acid soil at Mangoplah, NSW, Australia (35.3532054°S, 147.2901734°E*, Experiment 5*). Symptoms of Mn^2+^ toxicity were visually scored as tolerant (0) and sensitive (1). Lines segregation for Mn^2+^ tolerance (in control accessions) were scored as intermediate (0.5). Experiment 6 involved the evaluation of 20 DH lines from the Darmor-*bzh*/Yudal population with different *BnMTP8* alleles for tolerance and sensitivity to Mn^2+^ toxicity. Acid soil was collected from the top 30 cm layers from the Mangoplah site and tested for pH. The soil had a pH of 4.3 in CaCl_2_, and Mn content was 198µM. The soil suitability was validated using a bioassay conducted in an environment-controlled growth chamber (Percival, USA) with a 250 µmol M^-2^ S^1^ photon flux density, 50% humidity and a 22°C/20°C (16/8h) day/night temperature regime.

### DNA isolation

Genomic DNA was isolated following a standard phenol/chloroform extraction method for whole-genome resequencing and targeted gene sequencing of the *BnMTP8.A09* gene using the Sanger-sequencing approach at the Australian Genomic Research Facility (http://www.agrf.com). Low-density DArTseq genotyping of F_2_ lines based on the genotyping-by-sequencing method (Raman et al., 2014).

### Whole Genome Resequencing and SNP Identification

We resequenced 326 accessions using the WGR approach at the commercial Novogene and Illumina HiSeqXTen services (BGI-Shenzhen, China). Clean paired-end reads were mapped to the *B. napus* reference genome sequence, version 4.1 of the Darmor-*bzh,* downloaded from the Genoscope website (https://www.genoscope.cns.fr/brassicanapus, Chalhoub *et al*., 2014) using BWA-MEM with default parameters (Li and Durbin, 2009). The SNPs for each line were identified using the pipeline of Sentieon DNAseq (v201711.05, https://www.sentieon.com). The filtering was accomplished with function, variant filtration in GATK (v3.4-46-gbc02625, https://soCDFware.broadinstitute.org/gatk/) using the parameters of QUAL<30, MQ<50 and QD<2. The extent of heterozygosity and the minor allele frequency (MAF) were calculated using VCFtools (v0.1.13). High-quality SNPs with MAF <0.05 and missing rate >0.9 were used for GWAS analysis.

### Population Structure, GWAS and candidate gene identification

The tagSNPs, extracted using plink (v1.9) (Purcell et al., 2007), were used as input files for phylip software (v3.697) (https://evolution.genetics.washington.edu/phylip.html) to construct the phylogenetic tree. The tree was visualized using the FigTree package (v1.4.4, http://tree.bio.ed.ac.uk/soCDFware/figtree/). The population structure was constructed using admixture software (v1.3.0, www.genetics.ucla.edu/software/admixture). We estimated LD in VCFtools, using high-quality SNPs. The LD between marker pairs was estimated using the correlation coefficient of the allelic frequencies (*r^2^*), considering all the possible allele combinations. The physical distance when the decay of linkage disequilibrium (LD) reached the half maximum (1/2 LD distance) was calculated using VCFtools (v0.1.13, http://vcftools.github.io)) in a GWAS population. We accounted for the population structure and the kinship matrix for GWAS. The latter was calculated using the Efficient Mixed-Model Association eXpedited (EMMAX)-kin method. GWAS was conducted using the EMMAX software with a linear mixed model (Kang et al., 2010), which corrects spurious associations due to population structure. The LD heatmap, Manhattan and Quantile-Quantile (Q-Q) plots were drawn in the R program (v4.0.5, https://www.r-project.org/). The candidate genes for GWAS loci were extracted based on physical positions of highly significant associated SNP ± ½ LD distance (kb).

### Selective sweep analyses

The SNP data sets with missing rates < 0.1 were used for selective sweep analysis. The fixation index (*F_ST_*) and the nucleotide diversity (π) were analyzed using the VCFtools package. We analyzed a cohort of 50 extreme accessions (25 tolerant accessions with a mean score of < 0.2 and 25 sensitive accessions with a mean score of ≥4). The t-test of significance was used to determine the differences between tolerant and sensitive cohorts. The threshold of *F_ST_* >0.19, which corresponded to the top 5% of sites, was used to identify the selective sweep.

### *BnMTP8.A09* allele mining in *B. napus* global germplasm

Allelic variation, as SNPs and InDELs, in the *BnMTP8.A09* gene was investigated in the published dataset of resequenced 2,311 *B. napus* accessions, representing 1,259 accessions from Asia, 929 accessions from Europe, 60 from North America, two from South America, 38 from Oceania and four from Africa. (Song et al., 2020, Tang et al., 2021, Wu et al., 2019, Lu et al., 2019). These accessions included three ecotypes, spring (354 accessions), winter (756 accessions) and semi-winter (1,122 accessions). All the data were obtained from BnIR (https://bnaomics.ocri-genomics.net/).

### Genetic architecture of Mn^2+^ tolerance

To describe the genetic architecture underlying Mn^2+^ tolerance, the reported QTL region related to the Mn^2+^ tolerance of *B. napus* was integrated with significant SNP associations identified in the GWAS panel. Their physical positions or QTL intervals were aligned to the Darmor-*bzh* reference genome sequence of *B. napus* and were drawn using the MapChart program.

### *BnMTP8.A09* gene expression and structural variation analyses

We investigated gene expression using tails, i.e., extreme 20 phenotypes (i.e., Mn^2+^ tolerant with 1-2 score and Mn^2+^ sensitive with 4-5 scores) of the DH population derived from the Darmor-*bzh*/Yudal and GWAS panel (Table 1). Twelve biological replicates (4 plants/replicate x 3 replications) from each treatment were taken within one hour during the light cycle at ten days in hydroponic. Cotyledonary leaves showing tolerant and sensitive phenotypes were pooled from four plants per replicate and flash-frozen in liquid nitrogen. cDNA synthesis and the relative expression of the *BnMTP8.A09* gene were calculated by the relative quantification method, as outlined previously (Raman et al., 2019). Gene-specific primers for the *BnMTP8.A09* used for the expression analysis are given in Table S3. We also obtained sequence information for *MTP8* paralogs from whole-genome resequencing data of the 326 canola accessions. Variation across the *MTP8* paralogs was extracted using the gene model information or manually identifying gene regions based on BLAT homology (Table S4). The physical positions of different *MTP8* paralogs (NCBI GenBank accession; <u>AT3G58060.1</u>, MTP8 cation efflux family protein) were confirmed with those of the sequenced *MTP8* genes on the ‘Darmor-*bzh*’’ assembly v4.1. For each accession, the *BnMTP8* nucleotide sequences were aligned using MUSCLE as implemented in the software package Geneious (https://www.geneious.com). Structural variation, the number of polymorphic sites within the gene and the promoter region were identified. The selection pressure (*K_a_*/*K_s_* values for paralogous genes in *Arabidopsis*) of target genes was calculated using KaKs_Calculator v3.0 software (Zhang, 2022). The functional domains were verified using information from the NCBI conserved domain database.

### eQTL mapping on *BnMTP8.A09*

To analyse the expression regulation and potential interaction of *BnMTP8.A09*, we collected the RNAseq data and genotyping data of *B. napus* population from the Databases, including BnIR (http://yanglab.hzau.edu.cn/BnIR) and National Genomics Data Center (GSA Bioprojects: PRJCA002835, PRJCA002836 and PRJCA013095). The gene expression level was determined by TPM (Transcripts Per Million) and then used for eQTL mapping on target genes. The normalized gene expression values were used as the phenotype for eQTL analysis. GWAS-SNPs with MAF > 0.01 were utilized to perform eQTL mapping using Genome-wide Efficient Mixed Model Association (Zhou and Stephens, 2012) to detect associations of SNP-gene pairs. The threshold value for determining significant associations was set as –log_10_(1/n), where ‘n’ represents the total number of SNPs in the *B. napus* population. Based on the distance between eQTL and target genes, we subdivided an eQTL into *cis-*eQTL if its lead eSNP was found within 1 Mb from the transcription start site or transcription end site of the target gene; otherwise, including located on different chromosomes, it was assumed as *trans-*eQTL.

### Functional complementation in yeast

We used the hypersensitive yeast strain *pmr*Δ for functional complementation assay (Experiment 6). The coding sequence of the *BnMTP8.A09* gene (*BnaA09g37250D*) was translated to a peptide sequence and then backtranslated using the EMBOSS backtranslate tool with *Saccharomyces cerevisiae* codon preferences. A gene cassette was designed *in-silico* by incorporating the *S. cerevisiae* Tef2 promoter (700bp) and *S. cerevisiae TDH1* terminator (224bp) flanking the *S. cerevisiae* codon-optimized coding sequence. The construct was synthesized (Azenta) with a *Not*I site 5’ of the promoter sequence and *BamHI* site 3’ of the terminator sequence and was cloned into a polylinker of the low-copy yeast expression vector PRS-413 (Euroscarf). The plasmid was transformed into yeast knockout collection strain BY4741 *ΔPMR1* by incubating exponential phase cells at 42°C in 360µl of 50% PEG 3500, 100mM Lithium Acetate, containing 10µl herring sperm (Sigma). Transformed cells were plated on *Saccharomyces cerevisiae* medium with histidine dropout (Sigma). Vector-only controls were transformed into BY4741 *ΔPMR1* and By4741. Cells were normalized to an optical density of 1, and a serial dilution was plated on yeast minimal media without histidine and with various concentrations of MnCl_2_. Spot assays were photographed after 48 hours.

## Acknowledgements

The authors thank Mr. Kevin McCrea for making the Mangoplah field available for germplasm evaluation, Ms. Helen Burn for organizing the field site; Nawar Shamaya and Hannah Roe for help in phenotyping; Warren Bartlett for a sowing field trial and Novogene Corporation Inc., China, for resequencing accessions.

## Authors contribution

HR conceived the project, experiments, and research plan and wrote the manuscript with contributions from SB, HG and BP. RD provided the DH population from Darmor-*bzh*/Yudal. HR, BM, HG RR, SM, NK and SB conducted research, and HR, ZB, RR, YZ, NK and SL analyzed the data. All authors read and approved the final version.

## Data availability

The data supporting the findings and supplementary data are available within the paper. Resequencing data is being submitted to NCBI.

## Supporting information

**Table S1.** Accessions used to assess natural variation in tolerance to manganese toxicity in *B. napus*.

**Table S2.** Genetic variation for tolerance to Mn^2+^ toxicity in the GWAS panel

**Table S3** Sequence coverage of accessions used for GWAS analysis.

**Table S4** Principal component analysis of 326 accessions used for genome-wide association analysis

**Table S5** Genome-wide association analysis of manganese tolerance in *Brassica napus* accessions

**Table S6** Selective sweep analysis of selected 50 accessions of *B. napus* which showed contrasting variation in Mn^2+^ tolerance

**Table S7** Segregation ratio of F_2_ populations derived from five crosses. Mn^2+^ tolerance was evaluated in a nutrient solution supplemented with 125 µM MnCl_2_ (pH4.5). *: Non-significant. –: Data not suited for Chi-squared test

**Table S8** Candidate genes associated with manganese tolerance to manganese toxicity in a genome-wide association panel of 326 *B. napus* accessions

**Table S9** *BnaMTP8* homeologous genes in the reference genomes (cv. Darmor-*bzh* and ZS1).

**Table S10.** Diverse accessions used for *BnMTP08.A09* gene expression using RT-PCR. T: Mn^2+^ tolerant and S: Mn^2+^ sensitive.

**Table S11** List of primers used for sequencing of *MTP8* gene on A09 and quantitative RT-PCR.

**Table S12**. eQTL associated with manganese tolerance in *B. napus*.

**Figure S1:** Selective sweep and physical mapping of GWAS-SNPs associated with Mn^2+^ tolerance. A: Selective sweep signals between 25 Mn^2+^ tolerant and 25 Mn^2+^ sensitive accessions of *B. napus.* The dashed line represents the thresholds (top 5% of FST values) between 25 Mn^2+^ tolerant (group 2) and 25 Mn^2+^ sensitive (group 1) accessions of *B. napus*. The physical locations of significantly associated markers on chromosome A09 (B), C03 (C) and C09 (D) with Mn^2+^ tolerance; the closest markers and candidate genes are in green and red, respectively. The position of whole genome resequencing-based markers (with WGR suffix) is given in base pairs. String analyses of candidate genes (MTP8, AT3G58060 on A09, E; TMN1 on C03, F and ABCC5 on C09, G) associated with Mn^2+^ tolerance in *B. napus* showing both physical interactions and functional associations between known and predicted proteins interactions.

**Figure S2.** Frequency distributions of manganese tolerance scores in the F_2_ populations derived from P3083/ZY003 (A), Mutu/RSO94-67, (B), Darmor/Mutu (C) and Darmor/Jet Neuf (D)). F_2_ lines were evaluated for Mn^2+^ tolerance in a nutrient solution (Raman et al 2017) supplemented with MnCl_2_ (125 µmolar, pH 4.5).

**Figure S3.** Physical localisation of significantly associated DArTseq markers with manganese tolerance in *B. napus* F_2_ population derived from A: P3083 (China, tolerant to Mn^2+^) × ZY003 (China, sensitive to Mn^2+^).

**Figure S4.** Phylogenetic and sequence analyses of the *BnMTP8.A09* gene. A: Sequence variation in the BnaA09g37250D (*BnMTP8.A09*, 1966 bp) gene encoding cation diffusion facilitator protein in Darmor-*bzh* (Mn^2+^ tolerant), Yudal (Mn^2+^ sensitive) and selected doubled haploid lines (10 tolerant: T and 10 sensitives, S) derived from Darmor-*bzh*/Yudal. Seven exons and six introns are shown in blue and black, respectively. B: Gene structure and population SNPs/InDels distribution of six *BnMTP8*. C: Phylogenetic analysis of *BnMTP8.A09* homologues in *B. napus* and its ancestral diploid species. The tree was drawn using the neighbour-joining method in Geneious for MTP8 amino acid sequences from *B. rapa*, *B. oleracea* and *B. napus*. Arabidopsis *MTP8* gene was used as an outgroup. *BnMTP8* homologue that showed significant association with Mn^2+^ tolerance is labelled red.

**Figure S5.** *BnMTP8.A09* expression difference in Darmor-*bzh* (Mn^2+^ tolerant), Yudal (Mn^2+^ sensitive) and selected doubled haploid lines (11 tolerant: T and 11 sensitives, S) derived from Darmor-*bzh*/Yudal population are presented in Table Sx; Primer-pair used for *BnMTP8.A09* gene sequencing is given in supplementary Table 11.

**Figure S6.** Micronutrient analysis of selected 19 diverse *B. napus* lines which showed contrasting phenotypes for Mn^2+^ tolerance in nutrient solution. Micronutrients were determined by Inductively coupled plasma atomic emission spectrometry, following Delhaize et al (2007) at the Charles Sturt University, Wagga Wagga, Australia.

**Figure S7.** Structural variants and their distribution in *BnMTP8.A09* (*BnaA09g37250D)* gene. A: Different tissue types used for RNA library construction. B: Expression patterns of *MTP8* homologues in different tissue types e variants in the population; C: The violin plot reveals the difference of gene expression level of *BnMTP8.A09* between the population allelic variation of target InDel (A09-26819418). C Dendrogram showing the grouping of InDeL AT/A among winter, semi-winter and spring lines. D: Frequency of InDel among different *B. napus* ecotypes. E: The violin plot reveals the difference in seed oil content between the population allelic variation of target InDel (A09-26819418) located in *BnMTP8.A09.* Box shows the median and interquartile range values. “n” is the accession number for statistics. Seed oil content data from (Tang et al., 2021). *p*-value:5.196e-05 (*t*-test); 0.0338 (Wilcoxon-test).

**Figure S8.** Experimental design of doubled haploid lines (DH) from Darmor-*bzh*/Yudal implemented at the Mangoplah field site, NSW, Australia (A) and climate data of 2022 season (B). The trial was sown on 25^th^ April and harvested in December.

**Figure S9.** Selective sweep analysis of *BnMTP8.A09* gene in a GWAS panel of *B. napus. Fst* (A) and Diversity: π (B) analyses were performed using *B. napus* cv. ZS11 reference assembly.

